# Cross-modal applications of a neuromorphic olfactory learning algorithm

**DOI:** 10.64898/2026.06.02.727939

**Authors:** Michael L. Helde, Alexander G. Dimitrov

## Abstract

We adapted an olfactory neuromorphic algorithm to image and sound recognition. To achieve this, we carried out specific preprocessing procedures that were tailored to each modality. For images, we used the NIST digits dataset directly. For sound, we used samples from the Google Speech Command dataset. A gammatone filter was applied to each to reduce the noise of the short audio sample and convert the temporal sound signal to a positional frequency signal. The single stimulus test algorithm was then modified to handle audio processing on extracted columns from a gammatone filter spectrogram obtained from the sound file. We also implemented PCA for all modalities, retaining around 90% of the variance. The results showed that over sequential “olfactory” gamma cycles, the algorithm successfully achieved one-shot online learning over the image and sound modalities as well. However, PCA representations did not attain high similarities to their corresponding templates for all three modalities.

## 1 Introduction

The human brain is a highly advanced organ that governs all aspects of human behavior, composed of a vast network of neurons and synapses. Its complexity and functionality make it a fascinating subject for scientific exploration, as it outperforms modern computers in tasks like online learning, sensory/motor control, and interacting with the environment. Even the most powerful supercomputers are still unable to match the efficiency and sophistication of the human brain’s information processing capabilities. This has led researchers to explore the possibility of implementing the brain’s operating principles into the design of computing machines, which would result in highly efficient and powerful computing systems that could simulate the brain and aid in the study and treatment of brain-related diseases.

Neuromorphic computing is a type of computing system that draws inspiration from the structure and function of neurons and the networks they form. These systems mimic the brain’s functions more accurately than traditional computing systems, as they are designed to be analogous to the brain. The term “neuromorphic” was first used by Carver Mead in the 1980s (Mead, 1990) to describe very large-scale integration (VLSI) systems containing electronic circuits based on the biological design of the nervous system. Today, the term encompasses a broader range of concepts, including computing systems that use digital processors to simulate neural models of computation, as well as machine learning techniques that emulate biologically relevant neural networks and learning mechanisms.

In recent years, neuromorphic systems have gained attention for their potential to enable real-time, low-power processing of high-dimensional data. Various neuromorphic platforms—including Intel’s Loihi (Davies et al., 2018) and IBM’s TrueNorth (Merolla et al., 2014)—have been developed to support spiking neural networks (SNNs), a key model class for brain-inspired computation. SNNs have demonstrated capabilities in tasks ranging from vision and audition to control (Tavanaei et al., 2019; Roy et al., 2019). Their underlying plasticity mechanisms, such as spike-timing dependent plasticity (STDP), have also been studied as effective learning rules in unsupervised or semi-supervised settings (Bi and Poo, 1998; Masquelier et al., 2009).

Neuromorphic computing shows great promise for disrupting current computational and AI architectures. However, it is still limited by the lack of effective algorithms. Looking at the circuit-level organization of biological neural systems, which employ different computational strategies, presents a unique opportunity to develop novel biomimetic algorithmic approaches. It also positions neuromorphic computing systems as another powerful tool for brain research. Imam and Cleland (2020) developed a simplified spiking network model based on the architecture and dynamics of the mammalian olfactory bulb, demonstrating that it supports rapid online learning and signal restoration of odor inputs occluded by contaminants. This model can enhance the understanding of mammalian olfaction and improve the performance of artificial chemosensory systems. The authors suggest that this framework is also applicable to general signal identification problems with high-dimensional patterns embedded in noisy backgrounds (Hopfield, 1995). In the study presented here, we applied this neuromorphic algorithm to image and sound recognition to further test that hypothesis. We also investigated whether the algorithm relied on aspects of the structure of natural stimuli by comparing it to PCA representations of the same stimuli. Remarkably, the results show that the algorithm achieves one-shot online learning over image and sound modalities even though it was originally modeled after biological olfaction, indicating that the biological system may have arrived at a general solution for some high-dimensional signal processing. However, operations on PCA representations did not attain high similarities to corresponding templates in the training set for all three modalities, suggesting that the algorithm exploits statistical structure present in natural stimuli that is disrupted by PCA’s decorrelation and remains unrecoverable through simple rescaling or dimensional adjustments.

## 2 Methods

### 2.1 Mammalian Olfactory Bulb-modeled algorithm

A bio-inspired algorithm was developed by Imam and Cleland (2020) to model signal discrimination by the External Plexiform Layer (EPL) of the mammalian olfactory bulb. In our work, we repurpose that algorithm to process sensory modalities outside of its original use for odors.

To summarize the details presented by Imam and Cleland (2020, section Network architecture and plasticity), the neuromorphic model’s architecture operates as follows: The sensor input is received by the apical dendrite (AD) of each mitral cell (MC), stimulating its corresponding soma (S). This activation of MC is then transmitted through its lateral dendrites (see Fig. 1: depicted in green) to excite the dendrites of granule cells (GC, represented in purple) through synaptic connections. These excitatory connections, indicated by circles, are sparse and not influenced by spatial proximity. Conversely, the spiking activity of GC is conveyed as inhibition onto the local MC within the same column. GC spiking also initiated excitatory synaptic plasticity, implemented through a heterosynaptic STDP rule: the weights of synapses mediating presynaptic spikes immediately preceding a postsynaptic spike are strengthened, and the weights of all other incoming synapses are weakened. According to the authors, that produces effectively a ‘k-winners take all’ learning rule.

**Figure 1.**
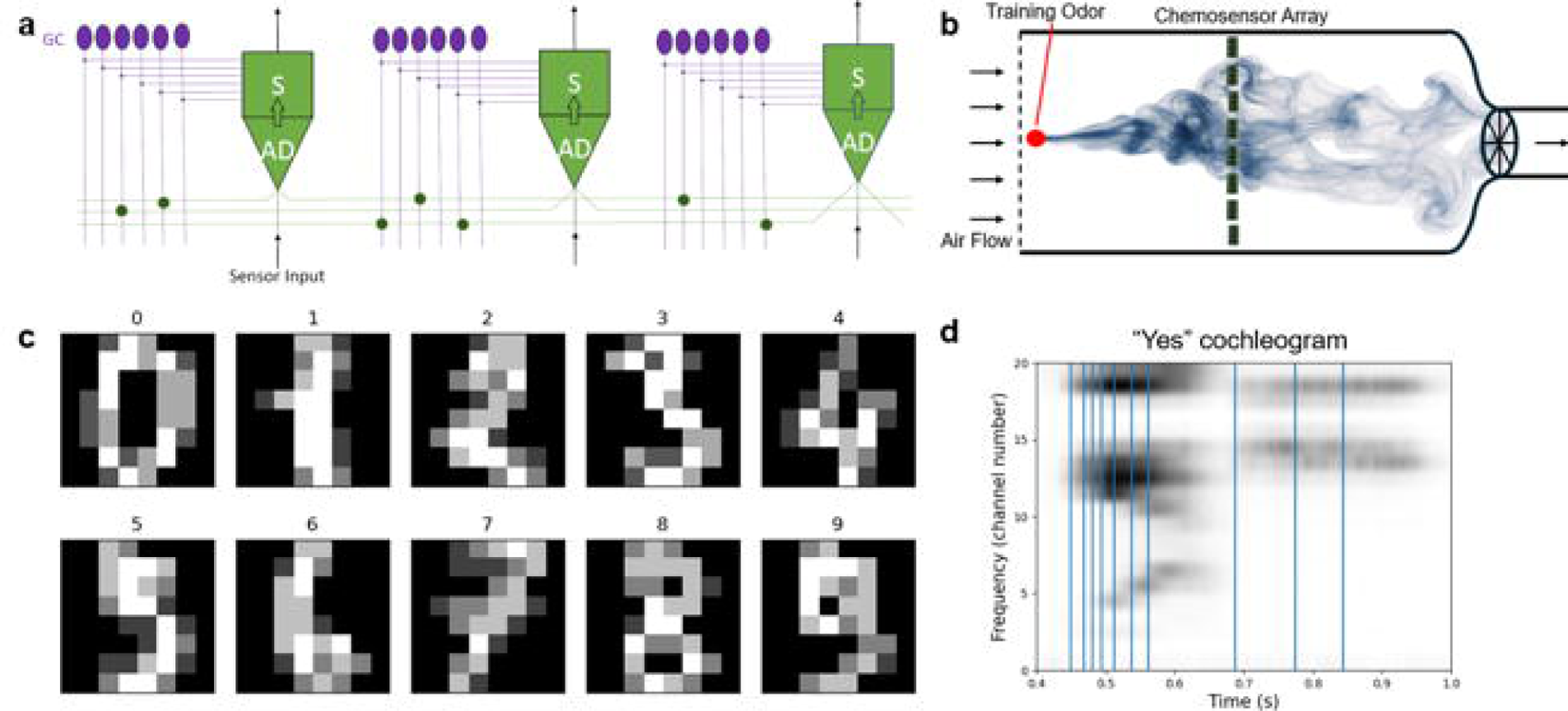
**(A)** A diagram of the neuromorphic spiking network inspired by models of the mammalian olfactory bulb (MOB), adapted from Imam and Cleland (2020), Fig. 1A. **(B-D)** The three modalities – the original olfactory Vergara et al. chemosensor array, the image NIST dataset, and “yes” audio cochleogram -- used in this paper to assess the generalizability of the neuromorphic algorithm. The odor data used here was nonoccluded presentations of odors without testing for variances in plume dynamics. Instead, for the multiple odor discrimination task, parametric occlusion was simulated allowing precise metrics of occlusion. An 8 × 8 pixelated representation of handwritten digits is used for assessing foveal-presented pattern recognition learning abilities. The vertical lines in panel d represent the manually selected time indices used for presentation to the algorithm. Each line is representative of 10 fine 20-dimensional channel number arrays spaced by 10 timesteps surrounding the central time index shown. See Methods sections 2.2 and 2.3 for preprocessing steps taken for images and audio, respectively.

The size of the network was matched to the size of the original dataset, which was determined by the 72 chemical sensors used to record odors. To match the dataset size for other modalities, the algorithm had to be generalized in order to work with stimuli of different sizes. The structure of the algorithm was also altered. The original algorithm interleaved one-shot learning and testing, as well as testing after full algorithmic neurogenesis. We modified the order to complete learning before evaluations on the training set, in order to assess the initial performance after full algorithmic neurogenesis. In both vision and audition cases, the initial learning stages were followed by an evaluation stage on a noise-modified test dataset. Algorithm performance was assessed on the test set. This assessment was taken using the Jaccard similarity coefficient which is the cardinality of the intersection of two sets divided by the cardinality of their union. Jaccard similarity is a useful metric for comparing sets regardless of their size.

The noise perturbation for test stimuli was also shared between modalities. In their work, Imam and Cleland (2020) use a model of olfactory occlusion noise: a random subset of measurements is selected and replaced with random numbers. Imam and Cleland (2020) use coarsely quantized stimulus representations (activity levels are integers between 0 and 15) based on the gamma cycle period of the network, resulting in sixteen distinct sensor activation levels. We preserve this design choice and quantize stimuli from all modalities to the same precision level. We then apply the original noise perturbation algorithm to simulate occlusion noise for all modalities.

### 2.2 Image Processing of Digits

We used the handwritten digits dataset imported by the scikit-learn machine learning library (Pedregosa et al., 2011). This dataset consists of 8×8 grayscale images of 10 handwritten digits (0-9, shown in Fig 1C) derived from the original NIST digit database through blockwise downsampling and quantization. Each image was represented as a 64-dimensional vector and quantized to integers in the [0, 15] range to match the input constraints of the EPL algorithm.

The NIST dataset was chosen for its compact and low-dimensional structure, which makes it suitable for testing biologically inspired algorithms not originally designed for detailed visual inputs. As the visual system is far more advanced in its organization than olfaction, the biological parallel to this discrimination task can be drawn to the ventral “object recognition” visual stream in humans and primates. Because the original EPL algorithm was designed for coarse olfactory data with 72 channels, NIST’s 64-pixel vectorization allowed for a more direct comparison without increasing the size of the network.

### 2.3 Auditory processing

We used the Google Speech Commands v0.02 dataset (Warden, 2018). Google’s sound database was used for monosyllabic words (e.g. “yes”, “no”, “up”, “down”). Shown in this paper is auditory analysis on “yes”. A gammatone filterbank was used to convert the auditory signal to a cochleogram (Slaney, 1998), simulating the initial stages of mammalian auditory processing. First, an array of center frequencies set in between the normal human speech and hearing frequency ranges (∼50 Hz - 17 kHz) (Pisanski et al., 2021; Purves et al., 2001) were constructed on the Equivalent Rectangular Bandwidth (ERB), scaled to continue the neuromorphic theme. Then, a bank of gammatone filters were implemented as cascades of four 2nd-order IIR filters. The filter bank size was varied from 10 to 100 to test performance on different stimulus sizes. The report is based on the results from the 20d filterbank. Lastly, the logarithm of the frequencies was taken mimic the nonlinear organization of the basilar membrane of the mammalian cochlea (Greenwood, 1990).

The approximately 1 second clip of a female speaker uttering the word “yes” was sampled at 16 kHz yielding fine timesteps associated with a fraction of a millisecond. Series of time index selections (the first two are 7200 and 7600 as shown in audio panels of Figures 2 and 3) were applied to the spectrograms generated of the short words to extract local patterns, within 100 steps of the one manually selected in steps of 10, suitable for input to the algorithm. Hence, each audio “clip” is presented to the algorithm by a 10 local samples of 20d frequencies. Time-step selections were made from a generated cochleogram of the word “yes” to sample across its three phonemes (Fig 1b). In accordance with the other modalities tested, the resultant vector was then quantized to have values in the range [0, 15] and used in the training and testing process of the EPL algorithm.

**Figure 2.**
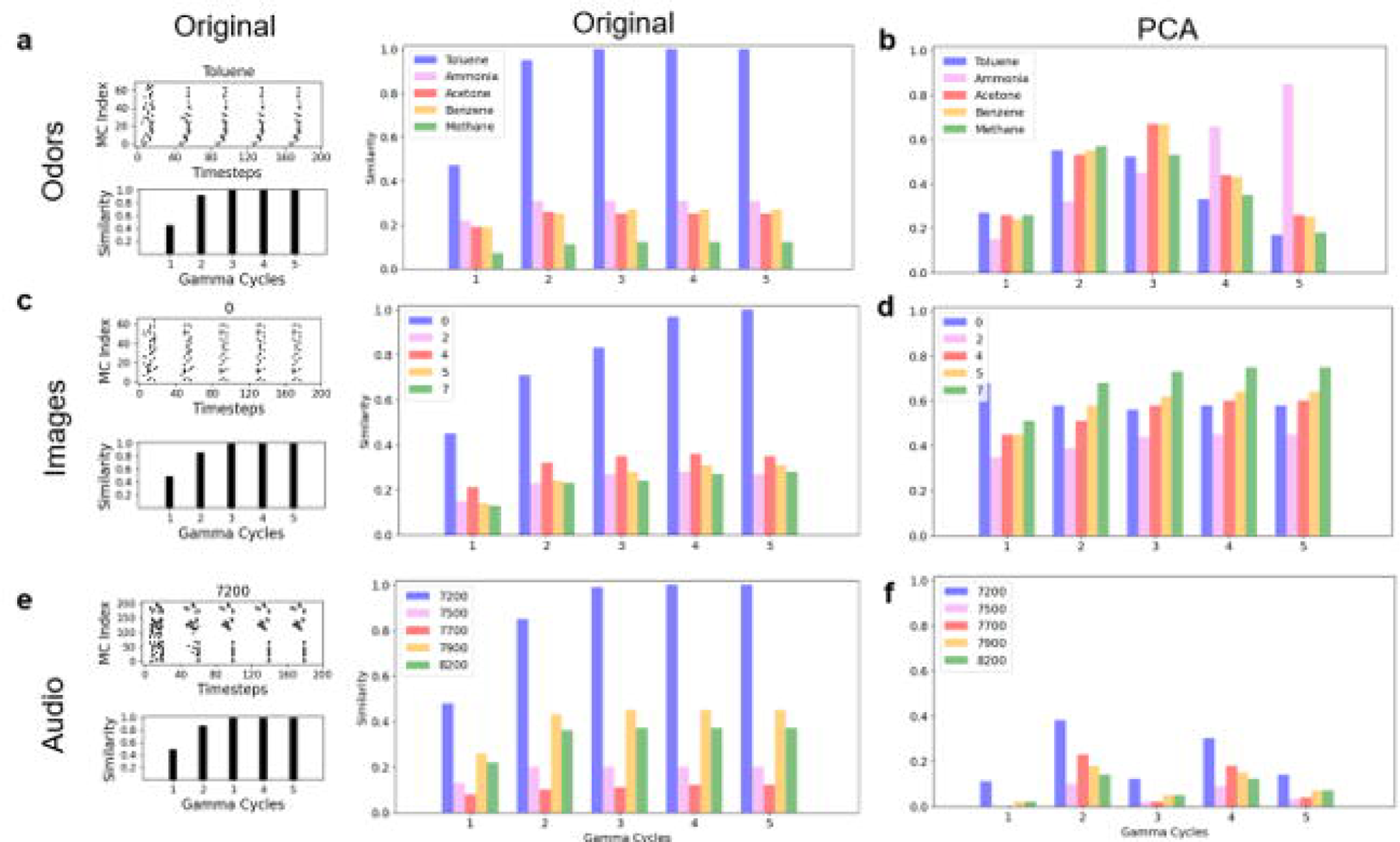
Multiple stimulus discrimination task performance across all tested modalities assessed via mitral cell raster plots and their associated Jaccard similarity index over successive gamma cycles. **(A, C, E).** Across the three modalities, an occlusion level of P=0.4 is used. To preserve clarity of summary plots, 5 of the 10 total stimuli are shown in the across-stimulus comparison plots for both the original stimulus set and their PCA counterparts **(B, D, F)**. Shown in blue is the similarity index between the learned representation of the training modal stimulus and the representation of the 40% occluded “testing” stimulus on its fifth gamma cycle presentation, or “sniff”. Notably, the algorithm can discriminate between the sample modal stimulus shown and the other learned stimuli. This is seen in mitral cell spiking activity by a sequential refinement of the spatiotemporal spike-train pattern.

**Figure 3.**
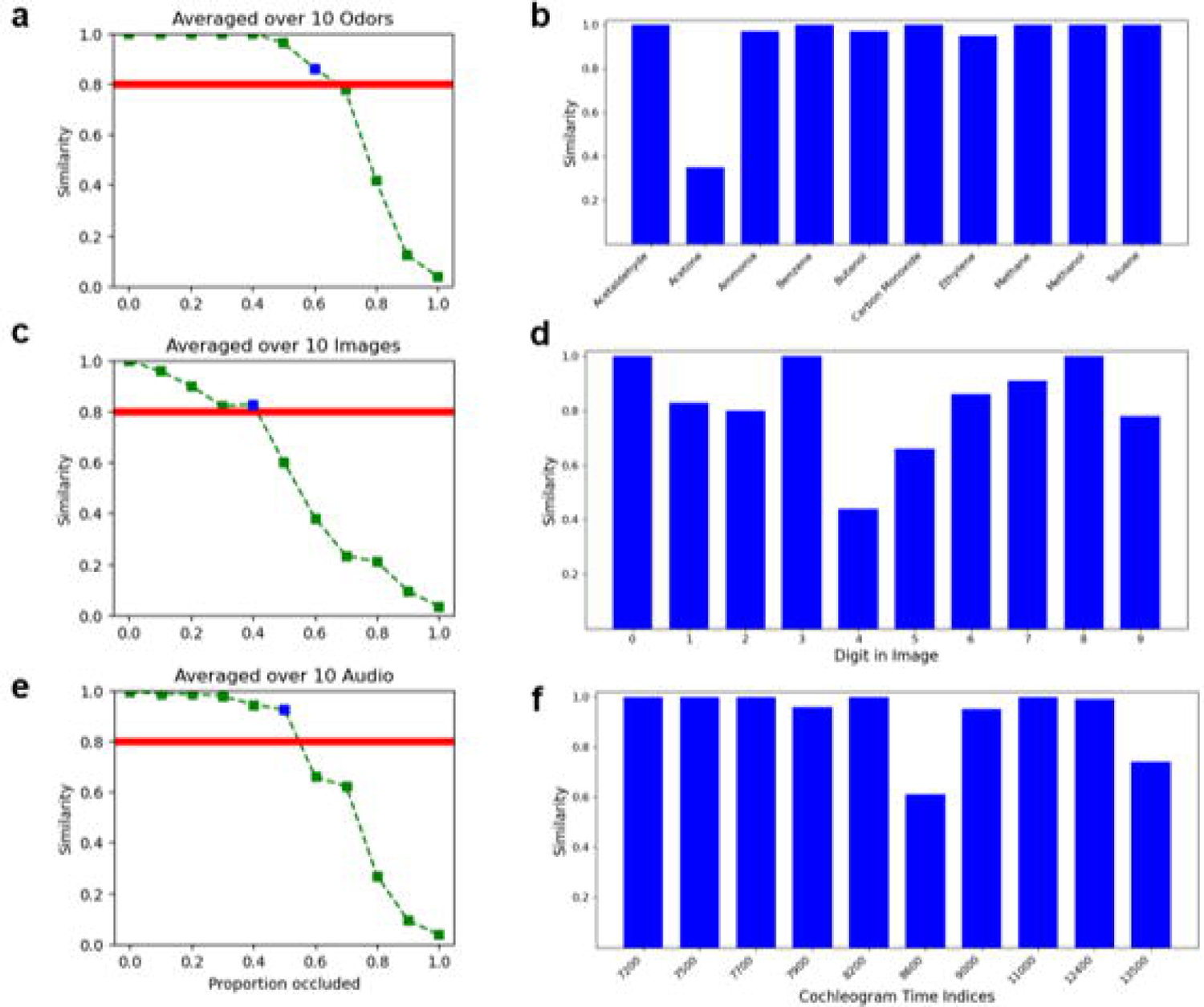
Ability of the algorithm to discriminate 10 stimuli across 11 occlusion levels (0 to 1 in steps of 0.1) on its highest-performing fifth gamma cycle. **(A, C, E)** The similarity between learned representations of stimuli and their code-generated occluded versions takes a roughly sigmoidal shape and crosses the 0.8 similarity threshold just after the occlusion level represented in blue. This occlusion level varies across modalities, and its associated performance across all stimuli over which the blue points in the first column of panels averages over **(B, D, F)**.

### 2.4 PCA compression

Principal Component Analysis (PCA) was used to compress the original sample of all modalities, including the original odors, using the PCA function from scikit-learn decomposition library. The number of components used was determined through trial and error to find the best-performing choice. PCA was applied as a part of the preprocessing steps for audio, odor, and visual signals, with the whiten parameter being tested as well. The results were reported based on pre-whitened PCA, as this maintained similar ranges for each principal component, which retained information after the coarse quantization to [0, 15]. Noise was included in the test performance after PCA compression as before. Even though the concept of “occlusion noise” of the quantized PCA representation may not be intuitive, it was kept to maintain consistency in the neuromorphic processing, except for the specific manipulations performed in this study, such as different modalities and PCA representation. The number of principal components was selected to approximately match the dimensionality of each modality (72 for odor, 64 for images, and 200 for audio). Varying the number of retained components around these values did not produce significant changes in algorithm performance, indicating that the observed effects are not strongly dependent on the exact dimensionality of the PCA representation.

We tested both partial and full PCA representations. Partial PCA involved reducing the number of components while preserving ∼90% of the variance, whereas full PCA simply involved a change of coordinates without dimensionality reduction. In both cases, the algorithm failed to recognize templates reliably. This failure likely stems from the nature of PCA outputs: they are dense and decorrelated, losing both spatial structure and sparsity. In contrast, the EPL algorithm functions as a sparse pattern matcher. When fed PCA representations, the distributed encoding and lack of structural alignment appear to obscure the features this network depends on. This mismatch likely explains the uniformly poor performance of PCA representations across all tested modalities.

## 3 Results

Our work shows that, while the network accommodates different sensory modalities well, it does not interact properly with standard compression tools like PCA.

In Figure 2 we show the similarity of a single test sample, standardized at an occlusion level of 40%, against the templates in the training set. That is evaluated for all modalities in original stimulus representations **(A, C, E**) and their respective PCA representations (**B, D, F**). One-shot learning is achieved in all modalities without PCA compression across the five gamma cycles (high and increasing similarity values, against lower similarity values for non-target templates). This is shown by the occluded ‘target’ test sample shown in blue across the modalities. That was toluene for odors, the digit “0” for images, and the gammatone cochleogram at time index 7200 (0.45s at a 16 kHz sampling rate) for sounds. Similar results were observed when using other test samples as a comparison for all modalities. This was not the case with PCA compression. All comparisons of occluded samples to trained templates yielded low similarity values (**B, D, F**).

The main result of the report is summarized in Figure 3. The figure shows network activity after one-shot learning caused by the occluded learned stimuli. It recapitulates the networks performance under different noise conditions (Figure 4e in Imam and Cleland) for all modalities investigated here. The Jaccard similarity between the occluded stimulus and the learned representation of the corresponding stimulus across five gamma cycles is plotted. The figure shows similarity to training templates for all test stimuli, for all three modalities, and in original and PCA representation. The results here reflect a 10% noise level (best case scenario), to demonstrate that the lack of discriminability for the PCA representation is due to the representation itself, and not the occlusion noise levels. Similar to the results by Imam and Cleland (2020), occlusion levels up to ∼30-40% marginally decreases the similarity values to trained templates for the original stimulus representation (**Fig 3A, C, E**). The responses to the PCA representation continued to be low, indicating no detection of similarity in that case.

Additional preprocessing steps were tested to improve PCA-based representations, including rescaling and equalization of principal component magnitudes prior to quantization. These modifications did not improve performance and, in some cases, further reduced similarity to learned templates, reinforcing the conclusion that the limitation arises from the structure of the PCA representation itself rather than simple scaling effects.

## 4 Discussion

The EPL algorithm was derived by Imam and Cleland (2020) from computational features of the mammalian olfactory system. It was interesting and instructive to address the question hypothesized by the authors - is their structure adapted specifically to olfactory signals, or does it contain elements allowing it to perform general signal discrimination. The results reported here are divergent. The system performed remarkably well across several sensory modalities, producing equally high similarity scores for visual and auditory stimuli.

However, we were surprised by its inability to distinguish the same signals when those were converted to PCA coordinate systems. The initial intent of the PCA transformation was to decrease the dimensionality of the input while preserving most of the signal information. But repeated applications of the algorithm, up to almost full-dimensional representation (a PCA coordinate change, without dimensionality reduction) highlighted this persistent unexpected feature. We conjecture that the EPL algorithm relies on natural signal statistics in its operations (Einhäuser and König, 2010; Schwartz and Simoncelli, 2001), which have many correlations, so no individual dimension is too important, and constrain the stimuli to a smaller sensory manifold. In contrast, by design PCA produces de-correlated representations, for which changes in each dimension lead to significant changes in the stimulus representation. We do not claim to prove this hypothesis in the report presented here but put it forward as a possible explanation of the observed neuromorphic system performance.

This lack of discriminability in PCA space was robust across variations in preprocessing choices, including the number of retained components and post-PCA normalization strategies, suggesting that the effect is intrinsic to the decorrelated structure of PCA representations. The concern around stimuli statistics aligns with recent discussions regarding evaluation methodology in neuromorphic olfactory models, where the use of similarity-based criteria has been noted to introduce potential ambiguities in assessing generalization performance (Dennler et al., 2024).

One direction for future work is to extend auditory preprocessing by incorporating finer temporal index selection and convolutional spectrogram representations. Such approaches may better preserve local structure in the signal and improve discrimination for repeated instances of the same spoken word. Another important direction is the implementation of this algorithm on neuromorphic hardware platforms such as Intel’s Loihi chip. This would enable direct evaluation of computational efficiency, energy consumption, and scalability as well in the context of non-olfactory modalities, providing a more complete assessment of the practical advantages of this biologically inspired approach.

## 6 Generative AI disclosure

ChatGPT was used to generate the plume of Fig 1B. It is an illustrative plume that does not capture the statistics of the odorant data used in this report.

## 7 Author Contributions

AD identified the neuromorphic theme and initial neuromorphic algorithm. MH modified the algorithm and performed poly-modal discrimination analysis. AD and MH discussed project progress and identified directions to pursue. AD and MH wrote and edited the manuscript.

## Acknowledgments

The authors wish to thank Dr. Nabil Imam for sharing olfactory sensory data and for helpful discussions throughout the project.

